# Combined inhibition of MDM2 and PARP lead to a synergistic anti-tumoral response in p53 wild-type rhabdomyosarcoma models

**DOI:** 10.1101/2024.12.03.626050

**Authors:** Guillem Pons, Patricia Zarzosa, Gabriel Gallo-Oller, Amelie Wenz, Maris Lapins, Lia García-Gilabert, Júlia Sansa-Girona, Natalia Navarro, Miguel F. Segura, Aroa Soriano, Gabriela Guillén, Raquel Hladun, Josep Sánchez de Toledo, Soledad Gallego, Jordi Carreras-Puigvert, Lucas Moreno, Josep Roma

## Abstract

Targeting MDM2-p53 interaction to enhance p53 activity represents a promising antitumoral strategy for p53 wild-type pediatric tumors, including soft-tissue sarcomas. However, results from early-phase clinical trials in hematological and solid cancers have shown limited efficacy of MDM2 inhibitors, underscoring the necessity to explore more effective combinations with other agents. In this study, we provide results from a 22-drug combination screening demonstrating the therapeutic potential of combining siremadlin (MDM2 inhibitor) and olaparib (PARP inhibitor) for treating p53 wild-type rhabdomyosarcoma (p53 WT RMS). Cell survival, cell death, and apoptosis analysis revealed synergistic effects of combining siremadlin plus olaparib *in vitro*. The combination of both drugs led to a significant increase in p53 activity, as evidenced by p53 accumulation and K382 acetylation, along with an increased expression of bona fide p53 targets. Furthermore, treatment with both drugs resulted in significant reduction of tumor growth and increased mice overall survival when compared to single treatments and control in cell line-derived orthotopic xenograft models. Overall, our study demonstrates, for the first time, the synergistic effect of combining siremadlin plus olaparib in inhibiting p53 WT RMS tumor growth *in vitro* and *in vivo*. Our findings support the potential to study the combination of both drugs in clinical trials and warrants further investigation in other tumors with similar molecular features.

## Introduction

Rhabdomyosarcoma (RMS) is the most common soft-tissue sarcoma in children and adolescents, originating from embryonic mesenchymal cells with the potential for myogenic differentiation^1^. This childhood sarcoma accounts for approximately 4% of all pediatric tumors and has a 5-year overall survival rate of 70% for localized cases, which declines to below 30% in metastatic cases and less than 20% at relapse^1–3^. Currently, risk-adapted treatment is based in the combination of surgery, chemotherapy and radiotherapy, with no novel agents been incorporated into frontline therapy for the past decades. RMS is classified into two main molecular subtypes: fusion-positive (FP-RMS) and fusion-negative (FN-RMS), each exhibiting distinct clinical behaviors, genetic alterations, and prognosis^4,5^. However, despite these differences, a critical intersection in their pathogenesis lies in the MDM2-p53 axis, which has emerged as a central node in RMS biology.

In FP-RMS, the *PAX3-FOXO1* and *PAX7-FOXO1* fusions act as key oncogenic drivers, fundamentally reprogramming the transcriptome of FP-RMS cells and promoting the expression of downstream target genes essential for tumorigenesis^6,7^. These fusions enhance the transcription of receptor tyrosine kinases (RTKs) and growth factors, leading to the hyperactivation of the PI3K-AKT and RAS-MAPK signaling pathways^7–11^. Similarly, in FN-RMS, these pathways are frequently altered due to genetic mutations in RTKs and components of both signaling pathways^4,12,13^. The activation of the PI3K-AKT and RAS-MAPK pathways results in the phosphorylation of MDM2 by AKT and ERK, leading to its overactivation, increased stability, and subsequent translocation to the nucleus^14–20^. MDM2 functions as the primary E3 ubiquitin ligase for p53, ubiquitinating it and mediating its transport from the nucleus to the cytoplasm, thereby promoting its degradation via the proteasome^21–23^. Consequently, the overactivation of MDM2 inhibits the tumor-suppressive functions of p53, allowing RMS cells to evade p53-mediated cell cycle arrest and apoptosis^24–26^. Thus, excessive suppression of p53 by MDM2 overactivation can diminish the efficacy of conventional treatments, including chemotherapy and radiation, which often rely on intact p53 pathways to induce tumor cell death^27,28^.

The oncogenic influence of PAX3-FOXO1 and PAX7-FOXO1 fusions on the MDM2-p53 axis is further enhanced by the significant upregulation of the transcription factor MYCN, which directly increases *MDM2* expression^29,30^. Additionally, genetic alterations such as *MDM2* amplifications, particularly prevalent in FN-RMS, and *CDKN2A* deletions, reinforce the role of MDM2 in RMS by providing additional mechanisms for p53 inhibition^4,12^. Notably, despite the higher mutational burden observed in relapsed/refractory (R/R) RMS compared to primary tumors, the majority of them retain the wild-type (WT) p53 status^31^. This persistence underscores the potential of MDM2 inhibitors to disrupt the interaction between MDM2 and p53 across both subtypes, presenting a novel strategy for developing targeted and effective therapies for this highly aggressive pediatric cancer.

In recent years, several small-molecule inhibitors that disrupt the interaction between MDM2 and p53 have emerged as promising strategies for treating p53 WT tumors. These inhibitors are able to block p53 degradation, increasing its levels, and potentially restoring its activity^32–34^. Activation of p53 through MDM2 inhibitors has been shown to inhibit the proliferation of p53 WT cells *in vitro* and *in vivo*^33,34^. However, the initial expectation that MDM2 inhibitors would activate p53 and induce significant apoptosis in p53 WT cells has been challenged by studies demonstrating that these compounds often result in limited apoptosis, largely due to specific patterns of p53 activation^33,35,36^. Furthermore, early-phase clinical trials have shown limited efficacy of MDM2 inhibitors as monotherapy, highlighting the need for combination therapies to enhance their antitumor effects while reducing dose-limiting toxicities and adverse effects associated with these compounds^37–39^.

Currently, multiple combination strategies involving MDM2 inhibitors are being explored in preclinical and clinical studies^33^. Given that MDM2 inhibitors are able to activate p53 WT without genotoxic effects, these compounds are strong candidates for enhance the therapeutic efficacy of existing chemotherapy and radiotherapy regimens while minimizing the risk of resistance associated with single-agent treatments^33,34^. Additionally, the combination of MDM2 inhibitors with other targeted therapies represents a promising strategy to overcome the mild-to-moderate therapeutic effects observed with many of these agents as monotherapies. Notably, simultaneous treatment of p53 WT cells with MDM2 inhibitors and apoptotic inducers, such as BCL-2 inhibitors, has shown effectiveness in preclinical settings, with idasanutlin and venetoclax currently under investigation in clinical trials^40–43^. MDM2 inhibitors are also being evaluated in combination with tyrosine kinase inhibitors and inhibitors targeting the MAPK and PI3K/AKT/mTOR pathways, given the interconnectivity of these commonly altered pathways in cancer^44–49^. Overall, the exploration of diverse combination regimens involving MDM2 inhibitors presents a dynamic and promising landscape for cancer treatment. However, it is noteworthy that the current focus on these combinations predominantly concerns adult cancers, with limited attention given to pediatric cancers. Thus, while progress is being made in understanding and implementing combination strategies in the adult cancer context, the need for parallel research in pediatric oncology remains a critical frontier^50^.

In this study, we aimed to evaluate the efficacy of various MDM2 inhibitors, both as monotherapies and in combination with other targeted therapies or chemotherapeutics, for the treatment of p53 WT RMS. Previous preclinical studies have demonstrated promising outcomes when MDM2 inhibitors are combined with standard chemotherapeutic agents in RMS^51,52^. However, the prevailing emphasis on particular combinations has constrained the exploration of potential synergistic effects with newly developed or currently tested compounds for refractory or relapsed RMS. To address this limitation, we conducted a screening of 22 drug combinations to identify synergistic interactions. Notably, we highlight the therapeutic potential of combining siremadlin with selected agents, particularly olaparib. Our in-depth analysis of the combined effects of siremadlin and olaparib on RMS tumor growth, both *in vitro* and *in vivo*, as well as its integration with currently approved treatments, provides valuable insights into innovative multidrug therapeutic strategies that may enhance treatment efficacy for p53 WT RMS and other solid tumors.

## Materials and methods

### Cell lines and cell culture

Six different RMS cell lines were used in this study: RH18, RH36, CW9019, RD, RH4, and RH28. All RMS cell lines were cultured in Minimum Essential Medium with Earle’s Salts (MEM, Biowest) supplemented with 10% fetal bovine serum (FBS, Biowest), 2 mM L-glutamine (Biowest), 1 mM sodium pyruvate (Biowest), 1X MEM non-essential aminoacids (Biowest), 100 U/mL penicillin, and 100 μg/mL streptomycin (Biowest). Cultures were maintained at 37 °C in a 5% CO2 atmosphere and were regularly tested for mycoplasma contamination. Additional information about the molecular and histological characteristics of the cell lines used in the study is summarized in the Supplementary Table S1.

### Cell survival analysis and drug screenings

Cell survival assays were performed by seeding RH18, RH36, CW9019, RH28, RH4, and RD in p96-well plates at the indicated number of cells/well (Supplementary Table S2). After 24 hours, compounds were diluted in medium before being added to the cells at the indicated concentrations (Supplementary Table S3). After the treatments, cells were stained with a 0.5% crystal violet, 20% ethanol solution to assess cell survival. Dried crystals were dissolved in 15% acetic acid and absorbance was measured at 590 nm using an Epoch microplate spectrophotometer (Biotek). Percentage of cell survival was calculated by normalizing the absorbance values of each condition to their respective control in each experiment. Dose-response curves and IC_50_ values were calculated using non-linear regression approximation implemented in GraphPad Prism 6.0 (GraphPad Software). Drug combination effects were analyzed using the Bliss independence model^53^ either implemented in the software Synergy Finder 2.0^54^.

### RNA isolation, retrotranscription and quantitative real-time PCR

Total RNA was extracted from samples using the RNeasy Mini Kit (Qiagen) following the manufacturer’s instructions and quantified using a Nanodrop 2000 Spectrophotometer (Thermo Fisher Scientific). For retrotranscription, 1 μg of RNA was mixed with 1 μg of random primers (Thermo Fisher Scientific) in nuclease-free water and heated at 70 °C for 5 minutes. Then, 5 μL of 5X M-MLV reaction buffer (Promega), 5 μL of a mixture of 10 mM dNTPs and 200 U of M-MLV reverse-transcriptase (Promega) were added to the mixture and incubated at 37 °C for 60 minutes. Real-time PCR was performed by mixing 0.5 μL cDNA with 5 μL 2X TaqMan Universal Master Mix (Thermo Fisher Scientific), 0.5 μL TaqMan assays (Thermo Fisher Scientific), and 4 μL nuclease-free water in PCR tubes. Then, each reaction mixture was transferred to the wells of a MicroAmp 384-well plate (Thermo Fisher Scientific). PCR reaction was performed using an ABI PRISM 7900HT real-time PCR system (Thermo Fisher Scientific). A 40-cycle PCR was performed to detect the gene expression of *CDKN1A* (Hs00355782_m1), *MDM2* (Hs01066930_m1), *FBXW7* (Hs00217794_m1), and *MDMX* (Hs00159092_m1). The housekeeping gene *TBP* (Hs00172424_m1) was used as an endogenous control. Relative quantification of mRNA levels was performed by the 2^(−ΔΔCT) method^55^.

### Western Blot

Pelleted cells were lysed by resuspension in RIPA lysis buffer (Thermo Fisher Scientific) supplemented with Halt Protease and Phosphatase Inhibitor Cocktail (Thermo Fisher Scientific). Tumor samples (∼5 mm diameter) were resuspended in 200 μL RIPA lysis buffer and homogenized using a Bead Ruptor 12 (Omni International) for one cycle (20 seconds, 5 m/s). Sample lysates were kept on ice for 20 minutes for cell lysis and then centrifuged at 13,300 rpm for 15 minutes at 4 °C to remove debris. Total protein concentration was measured using the DC Protein Assay Kit (Bio-Rad). For electrophoresis, 20 μg of each protein sample were mixed with Laemmli loading buffer (Sigma-Aldrich), heated at 70 °C for 10 minutes, and loaded onto 10% or 12% SDS-polyacrylamide gels (SDS-PAGE) in a mini-PROTEAN Tetra electrophoresis cell (Bio-Rad). An electric field (35 mA per gel) was applied for protein separation. Proteins were then transferred onto a methanol-activated PVDF membrane (Cytiva) using a Mini Trans-Blot Transfer Cell (Bio-Rad) at a constant 200 mA and 4 °C for 2.5 hours in cold transfer buffer. Membranes were blocked for 1 hour in 5% non-fat dried milk or 5% BSA diluted in TBS-T and incubated overnight with primary antibodies. The next day, membranes were washed with TBS-T and incubated for 1 hour with HRP-conjugated secondary antibodies. After additional washes, membranes were incubated with Amersham ECL WB Detection Reagent (Cytiva) and exposed to SuperRX Fuji Medical X-ray films (Fujifilm). Exposed films were marked and digitized by scanning. Densitometric quantification of protein bands was performed using ImageJ^56,57^. A list of primary and secondary antibodies used in the study is available in Supplementary Table S4.

### Cell painting

Cell painting experiments were conducted according to the protocol described by Bray *et al.*^58^ with minor modifications. After 48 hours of chemical exposure, cells were washed with PBS, leaving 10 μl in the wells to minimize disturbance. MitoTracker (Thermo Fisher Scientific) was then added to a final concentration of 900 nM, and the cells were incubated for 30 minutes at 37 °C. Following the removal of MitoTracker, cells were fixed with 4% paraformaldehyde (Histolab) for 20 minutes. After fixation, cells were permeabilized with 0.1% Triton X-100 (Cytiva) for 20 minutes and stained with a solution containing Hoechst 33342, Wheat Germ Agglutinin, Phalloidin, SYTO 14, and Concanavalin A (all from Thermo Fisher Scientific) for 20 minutes. Finally, plates were washed, sealed, and stored at 4 °C, protected from light until image acquisition. Fluorescence microscopy was performed using a high-throughput ImageXpress Micro XLS (Molecular Devices), capturing six sites per well across five fluorescence channels for various cellular compartments: DNA (Hoechst), mitochondria (MitoTracker), Golgi apparatus and plasma membrane (Wheat Germ Agglutinin), F-actin (Phalloidin), nucleoli and cytoplasmic RNA (SYTO 14), and the endoplasmic reticulum (Concanavalin A). Excitation and emission wavelengths were optimized for each dye. Image processing and analysis were conducted with CellProfiler^59^ and CellPose^60^, including quality control, illumination correction, segmentation, and feature extraction. A total of 2,330 features were extracted from each cell and exported for multivariate analysis using Python 3.

### Cell death assay

RMS cells plated in 24-well plates and treated previously at the indicated experimental conditions were stained with 5 μg/mL Hoechst 33342 and 3 μg/mL propidium iodide (PI), added directly to the medium. Stained cells were incubated for 15 minutes at 37 °C, and stained nuclei were observed and photographed using a fluorescent microscope (λ_EX_/λ_EM_ = 350/461 nM) and counted using ImageJ^61^.

### Apoptosis assay

Cells were harvested from cell culture dishes at the indicated experimental conditions and resuspended in 100 μL Annexin V Binding buffer (BD Biosciences) at a density of 2 × 10^5^ cells/mL. Then, 100 μL APC Annexin V were added and samples were incubated for 15 minutes at room temperature (∼ 25°C) before adding 400 μL Annexin V Binding buffer and 0.5 μL Sytox™ Blue Dead Cell Stain. Samples were subsequently incubated at room temperature (∼ 25°C) for another 30 minutes prior to the measure of APC Annexin V and Sytox Blue Dead Cell Stain fluorescence intensities on single cells using a BD LSR Fortessa flow cytometer (BD Biosciences). Unstained cells and cells stained either with APC Annexin V or Sytox Blue Dead Cell Stain were used to adjust the live, apoptotic and dead populations. Cells treated with 10 nM actinomycin D (MedChemExpress) for 24 hours were used as a positive control of apoptosis. Flow cytometry results were analyzed using the FlowJo v10.8 Software (BD Biosciences). Cells negative for both markers were considered alive cells, Annexin V-positive cells were considered early apoptotic cells, Sytox Blue-positive cells were considered dead cells and cells positive for both markers were considered late apoptotic cells.

### Cell cycle analysis

Cells were harvested from cell culture dishes or plates at the indicated experimental conditions, resuspended in 300 μL PBS, and fixed at a density of 1 × 10^6^ cells/mL in 700 μL ice-cold ethanol overnight at −20 °C. After 24 hours, fixed cells were centrifuged at 5000 rpm for 5 minutes and washed twice with PBS to remove the fixing solution. Then, cells were incubated in a staining solution containing 0.1% Triton X-100 (Cytiva) and 10 μg/mL DAPI (Thermo Fisher Scientific) in PBS to a final volume of 1 mL. Cells were incubated at room temperature (∼ 25 °C) in the staining solution for 30 minutes prior to the measure of the DAPI intensity on single cells using a BD LSRFortessa flow cytometer (BD Biosciences). Flow cytometry results were analyzed using the Watson cell cycle univariate model ^62^ implemented in the FlowJo v10.8 Software (BD Biosciences).

### Cell-derived orthotopic xenografts

RMS cell-derived orthotopic xenografts (CDOX) were established by injecting 2 × 10^6^ RH36 cells into the gastrocnemius muscle of 5-week-old SCID mice (Charles River Laboratories). Tumor volume was monitored by measuring limb dimensions with a caliper and calculated using the following formula:

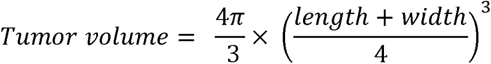

Mice were randomized and treated once the tumors were confirmed at approximately 25 mm^3^. In the pilot study, mice were randomized into 6 groups (*n* = 2) and received the following treatments: control (vehicle), 50 mg/kg siremadlin, 100 mg/kg olaparib, 50 mg/kg siremadlin plus olaparib 100 mg/kg, 25 mg/kg siremadlin plus 50 mg/kg olaparib, and 12.5 mg/kg siremadlin plus 25 mg/kg olaparib. In the experimental study, mice were randomized into 4 groups (*n* = 6): control (vehicle), 50 mg/kg siremadlin, 100 mg/kg olaparib, and 25 mg/kg Siremadlin plus 50 mg/kg olaparib). Both drugs were dissolved in a vehicle consisting of a mixture of 5% DMSO, 40% PEG300, 5% Tween-80, and 50% saline, added sequentially. In both studies, treatments were administered orally twice weekly, and animals were euthanized when tumor volume reached 1500 mm^3^. Ethical endpoint criteria, including acute weight loss (>10% of total body weight) or poor general appearance, were considered during the procedures. Tumor samples were collected immediately after euthanasia.

Simultaneously, an additional experiment was conducted to assess the molecular effects of the treatments following three dosage regimens (1, 3, and 5 doses). Mice were randomized into the same treatment groups as in the experimental study, but also according to the number of doses administered, resulting in 12 experimental conditions (*n* = 2). To ensure sufficient tumor sample size for subsequent studies, treatment was initiated when tumors reached approximately 250 mm³, with samples collected 24 hours after the last dose administration. All experimental procedures were conducted at the Rodent Platform of the Laboratory Animal Service of the Vall d’Hebron Research Institute (LAS, VHIR, Barcelona, Spain), under pathogen-free conditions. These procedures received prior approval from the regional Institutional Animal Care and Ethics Committee of Animal Experimentation of the Vall d’Hebron Research Institute (CEEA 70/19) and were performed in compliance with EU directive 2010/63/EU.

### Statistical analysis

Statistical analysis was performed using GraphPad Prism 6.0 (GraphPad Software). Parametric or non-parametric test were chosen based on the assessment of normality and equal variance of the data. Statistical significance between two groups were conducted using Student’s *t*-test or the Mann-Whitney *U* test. Statistical significance between three or more groups was determined by using one- or two-way ANOVA followed by Dunnett’s or Tukey’s *post-hoc* test for multiple comparisons. Each *p*-value was adjusted to account for multiple comparisons. The level of significance was denoted using asterisks (*) or hashes (#) as follows: *p* < 0.05 (* or #), *p* < 0.01 (** or ##), and *p* < 0.001 (*** or ###). Plots representing the data indicate the mean ± standard error of the mean (SEM) of three independent replicates, unless otherwise stated. Additional details regarding statistical analysis can be found on each section.

## Results

### p53 WT RMS cells were especially vulnerable to siremadlin

To investigate the differential drug responsiveness of p53 wild-type (p53 WT) and p53 mutant (p53 MUT) RMS cells, a single-drug screening was conducted. This screening included 26 distinct pharmaceutical agents, tested at three different concentrations across six RMS cell lines (three p53 WT (RH18, RH36, CW9019) vs three p53 MUT (RH28, RH4, RD)) (Figure S1a). The targeted therapies were selected for their specific relevance to pathways or proteins directly implicated in the p53 pathway. Moreover, first-line and second-line chemotherapies commonly used in the treatment of pediatric rhabdomyosarcoma were also included. Results from the screening revealed increased vulnerability of p53 WT RMS cells to the tested MDM2 inhibitors (siremadlin, idasanutlin, navtemadlin, and MI-773), with no significant differences observed for the other drugs (Figures 1a-b). To assess the potency and selectivity of the MDM2 inhibitors, IC_50_ values were determined for each compound. Siremadlin and idasanutlin exhibited lower IC_50_ values in p53 WT cells compared to navtemadlin and MI-773, indicating a greater potency of these compounds in inhibiting the MDM2-p53 interaction (Figures 1c and S1b-c). Notably, the difference in logIC_50_ values (ΔlogIC_50_) between p53 WT and p53 MUT cells was significantly greater for siremadlin, suggesting an enhanced selectivity of this compound for p53 WT cells over p53 MUT cells compared to the other MDM2 inhibitors (Figure 1c).

**Figure 1.**
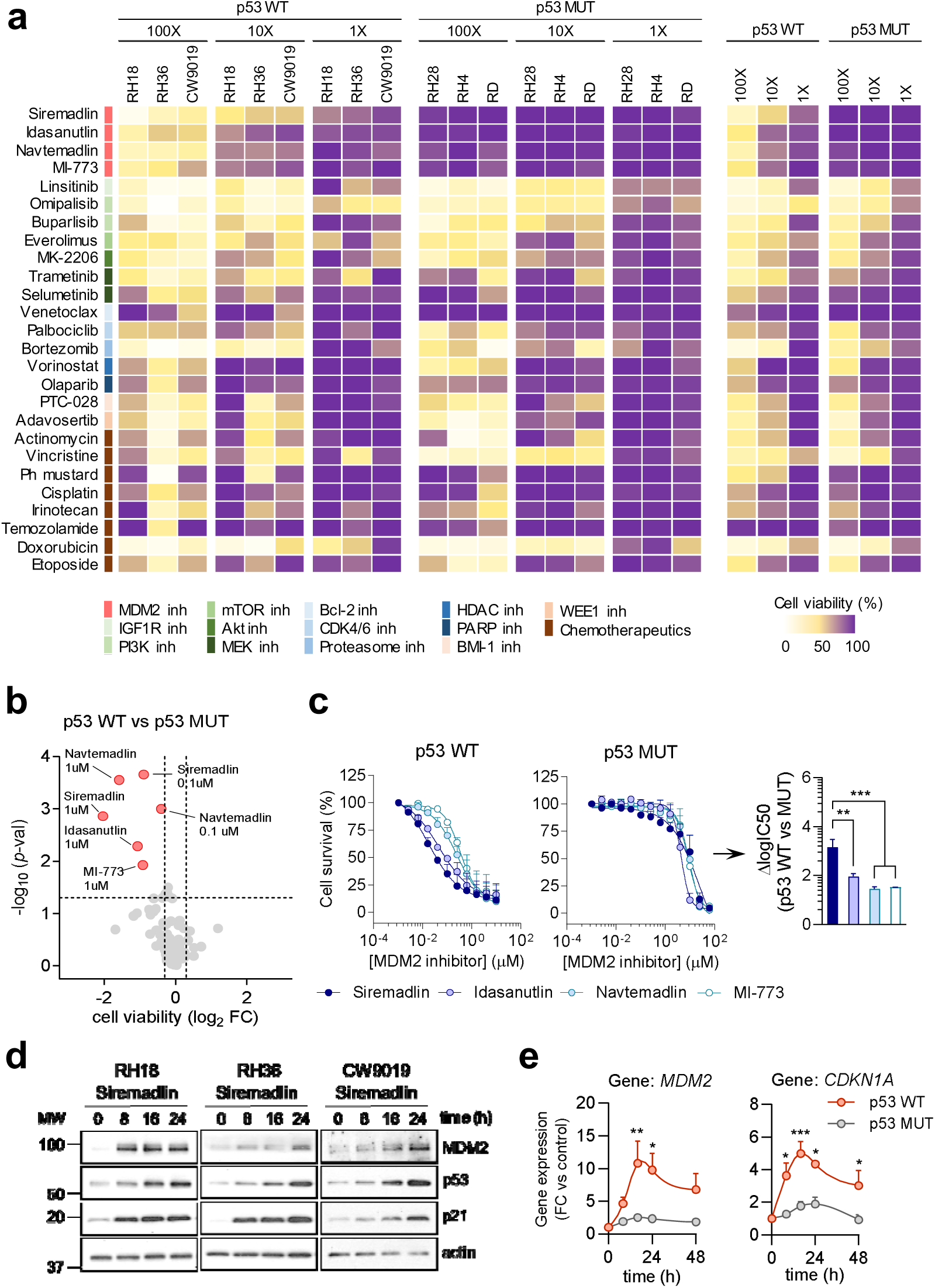
MDM2 inhibitors are able to reduce cell survival and modulate p53 activity in p53 WT RMS cells. (**a**) Heatmap summarizing cell viability results (in %) for 26 drugs tested at three concentrations (100X, 10X and 1X) in six RMS cell lines: three p53 WT (RH18, RH36, CW9019) and three p53 MUT (RH28, RH4, RD) cells, after 48 hours (*n* = 3). (**b**) Volcano plot showing the differential response of p53 WT and p53 MUT cell lines to the 26 drugs tested. The x-axis represents the difference in cell viability, expressed as log_2_ fold-change (log_2_FC), between p53 WT and p53 MUT cell lines for each drug and concentration, while the y-axis shows the associated significance level, expressed as -log_10_ p-value. Statistical significance was determined using Student’s t-test. Results were considered significant according to the following cut-off levels: p-value < 0.05, and abs(log_2_FC) > 0.25. (**c**) Dose-response curves of MDM2 inhibitors (siremadlin, idasanutlin, navtemadlin, and MI-733) are shown for p53 WT (left) or p53 MUT (middle) cell lines (*n* = 3). IC_50_ values were calculated using a non-linear regression approach (least squares regression without weighting). The right panel displays the difference in IC_50_ values between p53 WT and p53 MUT cells, expressed as the mean ± SEM. Statistical significance was assessed using one-way ANOVA followed by Tukey’s post-hoc test. (**d**) Western blot showing the expression of MDM2, p53 and p21 in p53 WT cells treated with 0.08 μM siremadlin for 0, 8, 16, and 24 hours. β-actin was used as loading control. Marks (dashes) on the left indicate the molecular weight (MW) markers on the membrane (**e**) Time-course plots showing the expression of MDM2 (left) and CDKN1A (right) in p53 WT (orange) and p53 MUT (grey) cell lines treated with 0.08 μM siremadlin for 8, 16, 24, and 48 hours (*n* = 3). Gene expression fold-change (mean FC ± SEM) is relative to untreated cells. Statistical significance was assessed using two-way ANOVA followed by Sidak’s post-hoc test.

Given that MDM2 and p53 expression may influence the sensitivity of RMS cells to MDM2 inhibitors, the expression levels of these proteins were examined in RMS cell lines, along with p21, a bona fide p53 target. No significant differences were observed in the protein levels of p53, MDM2, and p21 between p53 WT and p53 MUT cells (Figures S1d-e). Similarly, no significant differences were observed in the mRNA levels of MDM2 and p21 (*CDKN1A*) between p53 WT and p53 MUT cells, likely due to high variability in their expression among p53 WT cells (Figure S1f). Subsequently, to understand the molecular effects of siremadlin on p53 activity we assessed the levels of these proteins over time in p53 WT cells. WB results demonstrate a progressive accumulation of p53, along with an increased expression of MDM2 and p21, in p53 WT RMS cells at 8, 16, and 24 hours post-treatment (Figures 1d and S1g). Consistently, gene expression analysis by RT-qPCR revealed an increased expression of MDM2 and p21 in p53 WT vs MUT RMS cell lines after treatment with siremadlin for 16 and 24 hours, suggesting a potential induction of p53 transcriptional activity specifically in p53 WT RMS cells following the treatment (Figure 1e). The enhanced sensitivity observed in p53 WT cells to siremadlin, along with an increased p53 transcriptional activity evidenced by the accumulation of MDM2 and p21 after treatment, revealed an effective modulation of p53 activity by siremadlin in p53 WT RMS *in vitro*.

### The combination of siremadlin and olaparib synergistically reduced cell survival and induced morphological alterations in p53 WT RMS cells

To explore potential synergies or additive effects of siremadlin with other treatments, a drug combination screen was performed by treating p53 WT RMS cell lines with siremadlin in combination with the drugs used in the single-drug screening (Figure 2a). Cell viability and Bliss score were used to assess the effectiveness and nature of drug-drug interactions (synergism, additivity or antagonism). Notably, the combination of siremadlin with olaparib exhibited the highest Bliss score along with a substantial reduction in cell viability, emerging as the most promising combination (Figure 2b). To validate the screening results, p53 WT RMS cells were treated with increasing concentrations of siremadlin and olaparib, both individually and in combination, following a 5 x 5 matrix approach. Results from SynergyFinder 3D δ-score plots revealed additive to synergistic effects between the two compounds at the tested doses (Figure 2c). These effects were consistently observed across nearly all scores at 3 days post-treatment, suggesting an enhanced therapeutic effect of the combination beyond individual treatments (Figure 2d). Quantitative analysis of cell proliferation over time also showed a significant reduction in cell numbers at 5, and 7 days after treatment with siremadlin and olaparib, surpassing the effects observed with single treatments (Figure 2e).

**Figure 2.**
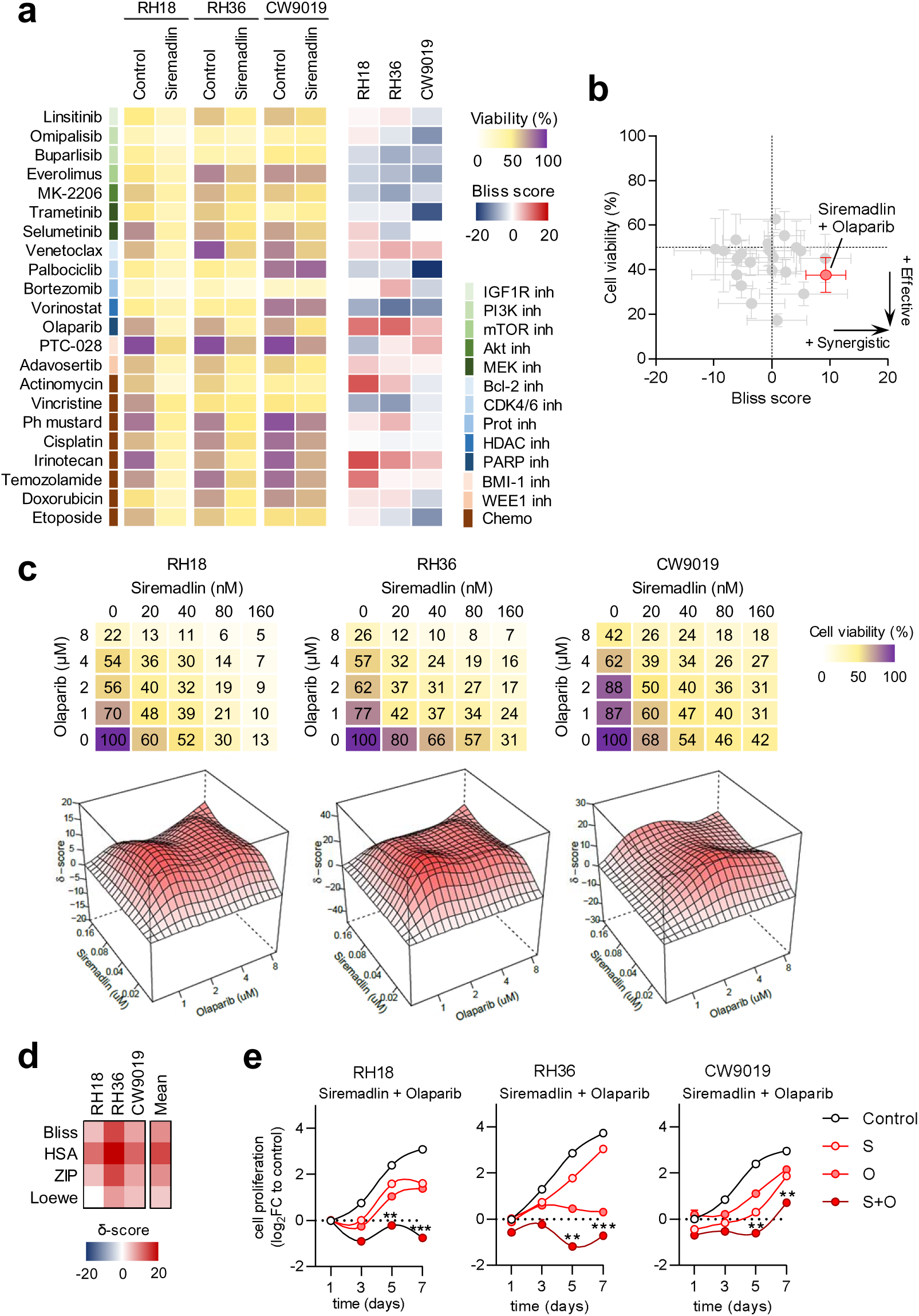
Synergistic antitumoral effects of combining siremadlin with olaparib in p53 WT RMS cells. (**a**) On the left, heatmap summarizing the effects on cell viability (in %) of p53 WT RMS cells treated siremadlin in combination with 22 different drugs for 48 hours. On the right, heatmap showing the degree of synergism (Bliss > 10), additivity (Bliss ≈ 0), or antagonism (Bliss < −10) between the drug combinations, based on the Bliss independence model. (**b**) Dot plot summarizing the results from panel (a). The excess over Bliss is displayed on *x*-axis, while the cell viability (in %) is displayed on *y*-axis. The lower-right quadrant highlights the most synergistic combinations with the greatest reduction in cell viability. Each dot represents the mean ± SEM from three p53 WT RMS cell lines (*n* = 3). (**c**) On the upper section, cell viability (in %) of p53 WT RMS cells treated with increasing concentrations of siremadlin (0–160 nM) and olaparib (0–8 μM) for 72 hours. Data represent the mean viability from three replicates (*n* = 3). On the lower section, SynergyFinder δ-score (Bliss score) plots generated from 5×5 dose-response matrices, indicating additive to synergistic effects between siremadlin and olaparib at the tested doses across each p53 WT RMS cell line. (**d**) Heatmap showing the mean synergy δ-scores for each cell line using four reference models: Bliss, Highest Single Agent (HSA), Loewe and Zero Interaction Potency (ZIP). The mean δ-scores across all three p53 WT RMS cell lines are summarized on the right. (**e**) Analysis of cell proliferation in p53 WT RMS cells treated with 80 nM siremadlin and 4 μM olaparib over 1, 3, 5, or 7 days. Cell proliferation is expressed as log_2_ fold-change (log_2_FC) relative to vehicle-treated control cells. Dots represent the mean ± SEM of three replicates (*n* = 3). Statistical significance was assessed using two-way ANOVA followed by Dunnett’s post-hoc test.

Building on the observed synergistic effects of siremadlin and olaparib on cell viability and proliferation, cell painting—a high-content, image-based assay that detects over 2000 morphological features—was used to investigate the phenotypic changes induced by the treatments and their impact on key cell structures and compartments (Figure S2a). Initial assessments compared the effects of siremadlin and olaparib alone with those exerted by compounds with known mechanisms of action (MoA). t-SNE analysis revealed distinct morphological clustering patterns, with olaparib and etoposide forming closely related clusters, suggesting a shared capacity to induce DNA damage in RMS cells *in vitro* (Figure S2b). In contrast, siremadlin produced a unique cluster, highlighting its distinctive morphological impact compared to other compounds with known MoA. Hierarchical clustering confirmed dose-dependent morphological changes for both drugs, with a heatmap illustrating the clustering of molecular profiles at increasing doses (Figure S2c). At high doses, siremadlin induced an increase in granularity across all measured cellular organelles, potentially indicating strong cellular stress, dysfunction, or the activation of pathways leading to cell death. In contrast, olaparib caused changes in the area and shape of the cytoplasm and nucleus, along with alterations in nuclear granularity, suggesting G2/M cell cycle arrest, where cells become enlarged after DNA replication in preparation for mitosis (Figure S2d). Remarkably, t-SNE analysis revealed a distinct clustering pattern for the combination of siremadlin and olaparib compared to each drug alone, indicating the induction of unique morphological changes in cells treated with the combination (Figure 3a). These changes were particularly evident in the ER, Golgi, cytoskeleton, nucleus, and intercellular measures (referred to as ‘neighbors’), such as the distance between adjacent cells, suggesting alterations in protein synthesis, trafficking, and nuclear processes, which may be linked to increased cell death induction (Figures 3b-c). Taken together, these results provide compelling evidence that the combination of siremadlin and olaparib synergistically induces morphological and functional changes beyond those of each drug alone. The combination potentiates alterations in multiple cellular compartments, underscoring its potential to disrupt essential cellular functions and highlighting its therapeutic relevance in targeting pathways involved in cancer cell survival.

**Figure 3.**
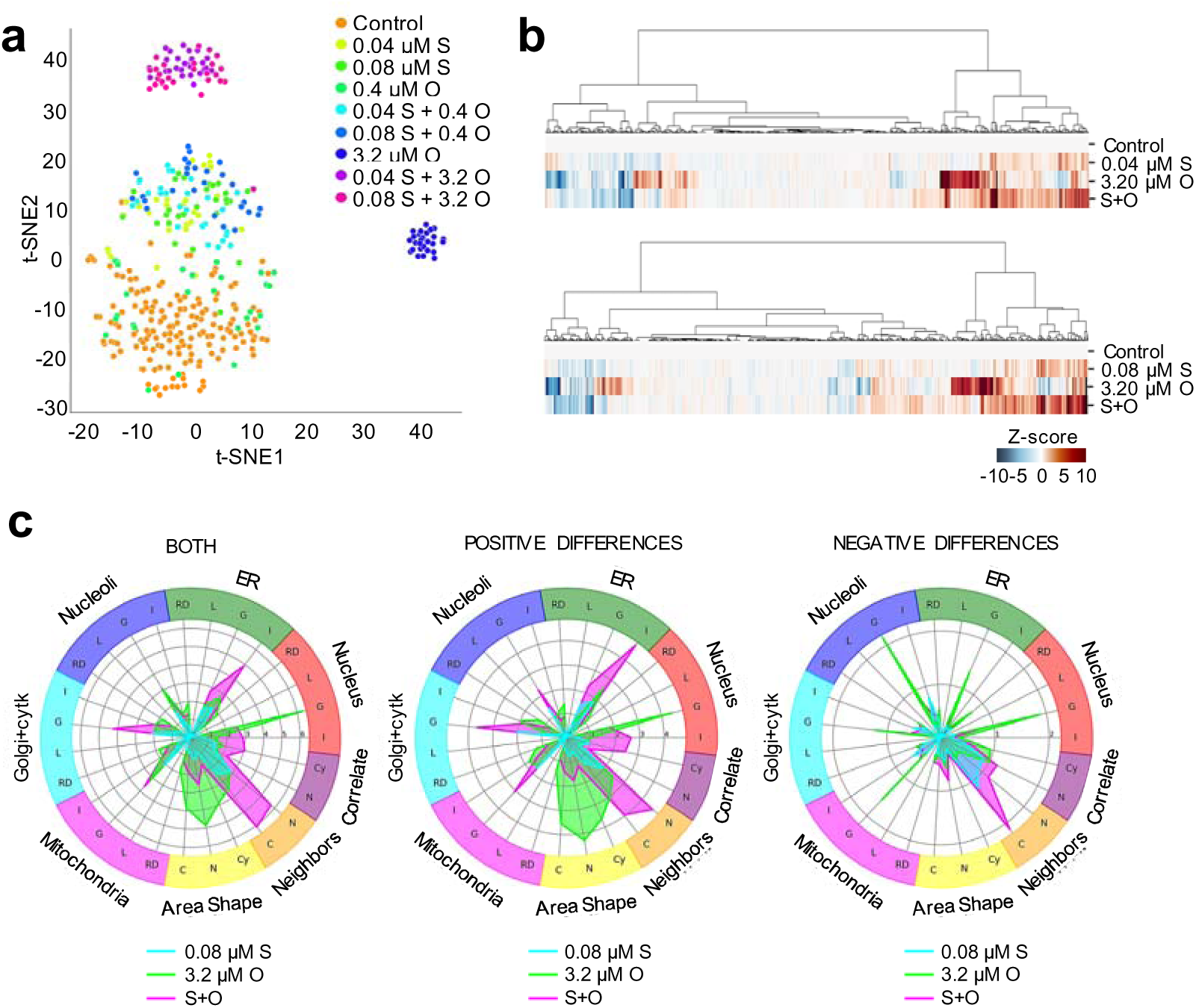
Combination of siremadlin and olaparib synergistically induce morphological changes in the ER, Golgi, cytoskeleton and nucleus. (**a**) Two-dimensional plot, derived from *t*-SNE analysis, showing the distinct morphological cluster formed by the combination of siremadlin (S) and olaparib (O) compared to individual treatments and control in RH36 cells. Scores on both axes represent the relative distance between clusters. (**b**) Heatmaps displaying the hierarchical clustering of >2000 morphological features analyzed by cell painting after treating RH36 cells with 0.04 or 0.08 μM siremadlin (S), 3.2 μM olaparib (O), and their combination (S+O). The color in the heatmap (Z-score) represents the deviation of each morphological feature from the mean value of the DMSO-treated control cells. (**c**) Radial plots showing the findings from panel (b) for RH36 cells treated with 0.08 μM siremadlin (S), 3.2 μM olaparib (O), and their combination (S+O). On the left, radial plots show the sum of both positive and negative differences compared to DMSO, while on the right results are categorized in positive and negative differences. Concentric marks represent Z-scores. All morphological features analyzed are grouped into eight main groups: endoplasmic reticulum (ER), nucleus, nucleoli, Golgi + cytoskeleton, mitochondria, cellular area and shape, characteristics of the neighbor cells, and correlates between nucleus and cytoplasm regions. At the same time, these features are further categorized into the following subgroups: RD (radial distribution), L (location and spatial distribution), G (granularity), I (integrated intensity), C (cell measurements), N (nuclear measurements), and Cy (cytoplasm measurements).

### The combination of siremadlin and olaparib synergistically induced apoptotic cell death in p53 WT RMS cells

Effective cancer therapies often involve the combination of agents that synergistically enhance cytotoxicity. In this study, the combination of siremadlin and olaparib demonstrated a synergistic reduction in p53 WT RMS cell survival and proliferation. To characterize the observed cytotoxic effects, we examined the combined effects of siremadlin and olaparib on cell cycle and cell death, using a set of complementary techniques.

Hoechst/PI double staining revealed a marked increase in cell death across all p53 WT RMS cell lines treated with the combination of siremadlin and olaparib for 72 hours, underscoring the potent cytotoxic effect of the combination compared to the single treatments (Figure 4a). Apoptosis was further assessed through Annexin V/Sytox staining followed by FACS analysis, unveiling a substantial induction of apoptosis in p53 WT RMS cell lines upon treatment with the combination (Figure 4b). These results suggested that siremadlin plus olaparib enhanced apoptotic cell death in p53 WT RMS cells, potentially through the activation of p53-mediated apoptotic pathways. Moreover, asynchronous cell cycle analysis using DAPI staining and flow cytometry revealed a G2/M cell cycle arrest in p53 WT RMS cells treated with either olaparib alone or the combination of siremadlin and olaparib (Figure 4c). To further elucidate the molecular mechanisms underlying these effects, WB was used to quantify the expression levels of phosphorylated retinoblastoma protein (pRb, Ser807/811) and Cyclin B1, key regulators of G_1_/S and G_2_/M cell cycle transitions. Notably, a substantial reduction in both pRb and Cyclin B1 levels was observed in p53 WT RMS cells treated with the combination for 72 hours (Figure 4d). The reduction of these markers suggested that the combination therapy interfered with both G_1_/S and G_2_/M checkpoints, thereby contributing to the previously observed antiproliferative effects. These findings highlighted the pharmacological effects of siremadlin and olaparib in combination therapy, demonstrating strong antiproliferative effects, increased apoptosis, and disrupted cell cycle progression in p53 WT RMS cells.

**Figure 4.**
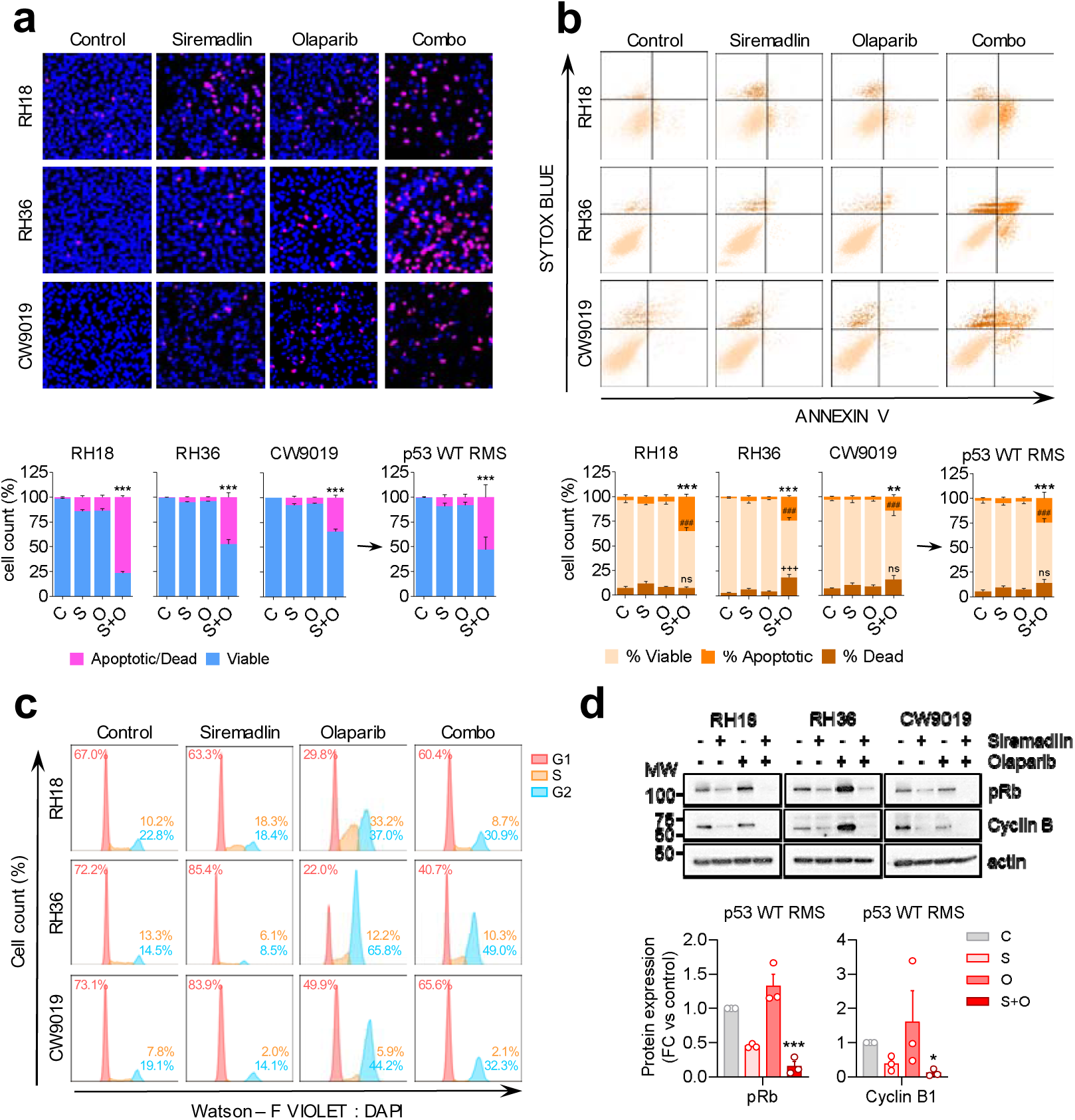
The combination of siremadlin and olaparib synergistically induces apoptotic cell death in p53 WT RMS cells. (**a**) On the upper section, images of Hoechst/PI double staining showing the effects on cell death induction in p53 WT RMS cells treated with 80 nM siremadlin (S), 4 μM olaparib (O), or their combination (S+O) for 72 hours. Viable cells are stained in blue while dead and apoptotic cells are stained in pink. On the lower section, bar plots representing the % of viable or dead/apoptotic cells for each cell line and treatment. Data are expressed as the mean ± SEM of three replicates (*n* = 3). (**b**) On the upper section, representative flow cytometry plots of Annexin V / Sytox double staining revealing the effects on apoptosis in p53 WT RMS cells treated with 80 nM siremadlin (S), 4 μM olaparib (O), or their combination (S+O) for 72 hours. On the lower section, bar plots representing the % of viable (Annexin V - / Sytox -), apoptotic (Annexin V +), or dead cells (Annexin V - / Sytox +) for each cell line and treatment condition. Bars represent the mean ± SEM of three replicates (*n* = 3). (**c**) Representative plots of DNA content from flow cytometry analysis of p53 WT RMS cells treated with 80 nM siremadlin (S), 4 μM olaparib (O), or their combination (S+O) for 72 hours. The area under each peak represents the number of cells in G1 (red), S (orange), or G2/M (blue) phases according to the intensity of DAPI staining. (**d**) On the upper section, WB showing the expression of p-Rb or Cyclin B in p53 WT RMS cells treated with 80 nM siremadlin (S), 4 μM olaparib (O), or their combination (S+O) for 72 hours. Molecular weight (MW) markers are indicated by dashes on the left side of the membrane. On the lower section, bar plots showing the quantification of p-Rb (left) or Cyclin B (right). Protein expression was normalized to β-actin and is expressed as fold-change (FC) relative to vehicle-treated cells. Data are presented as the mean ± SEM of three p53 WT RMS cells (*n* = 3). Statistical significance was assessed using two-way ANOVA followed by Dunnett’s post-hoc test for all experiments.

### The combination of siremadlin and olaparib increased MDM2 cleavage and p53 activity in p53 WT RMS cells

To further investigate the molecular effects of siremadlin and olaparib, the expression levels of PARP, MDM2, total p53, and acetylated p53 (ac-p53; K382) were quantified at 72 and 120 hours post-treatment. Additionally, the expression of p21 and BAX, which are targets of p53, were also evaluated to provide a comprehensive understanding of the molecular changes induced by the combination treatment.

In RH18 cells, siremadlin alone or in combination with olaparib reduced PARP expression, whereas olaparib alone led to a slight increase in both cleaved and total PARP at 72 hours (Figure 5a). In contrast, no significant differences in PARP or its cleaved form were observed in RH36 and CW9019 cells (Figures 5b-c). Notably, the combination of siremadlin and olaparib significantly increased MDM2 cleavage across all cell lines, suggesting enhanced caspase activity (Figures 5a-c and S3a). Consistent with MDM2 cleavage, p53 accumulation was observed in all cell lines treated with the drug combination at 72 hours (Figures 5a-c and S3b). This correlation suggested that MDM2 cleavage may contribute to p53 stabilization and/or accumulation in RMS cells. Additionally, increased levels of ac-p53 strongly suggested an enhanced p53 transcriptional activity (Figures 5a-c and S3b). In accordance with earlier findings regarding cell cycle progression, increased levels of p21 were observed in all cell lines at 72 hours following the combination treatment (Figures 5a-c and S3c). However, no significant differences were observed in BAX expression, suggesting that other proapoptotic proteins or molecular processes, such as the transcription-independent activation of BAX by p53, may contribute to the apoptotic process triggered by the combination (Figures 5a-c).

**Figure 5.**
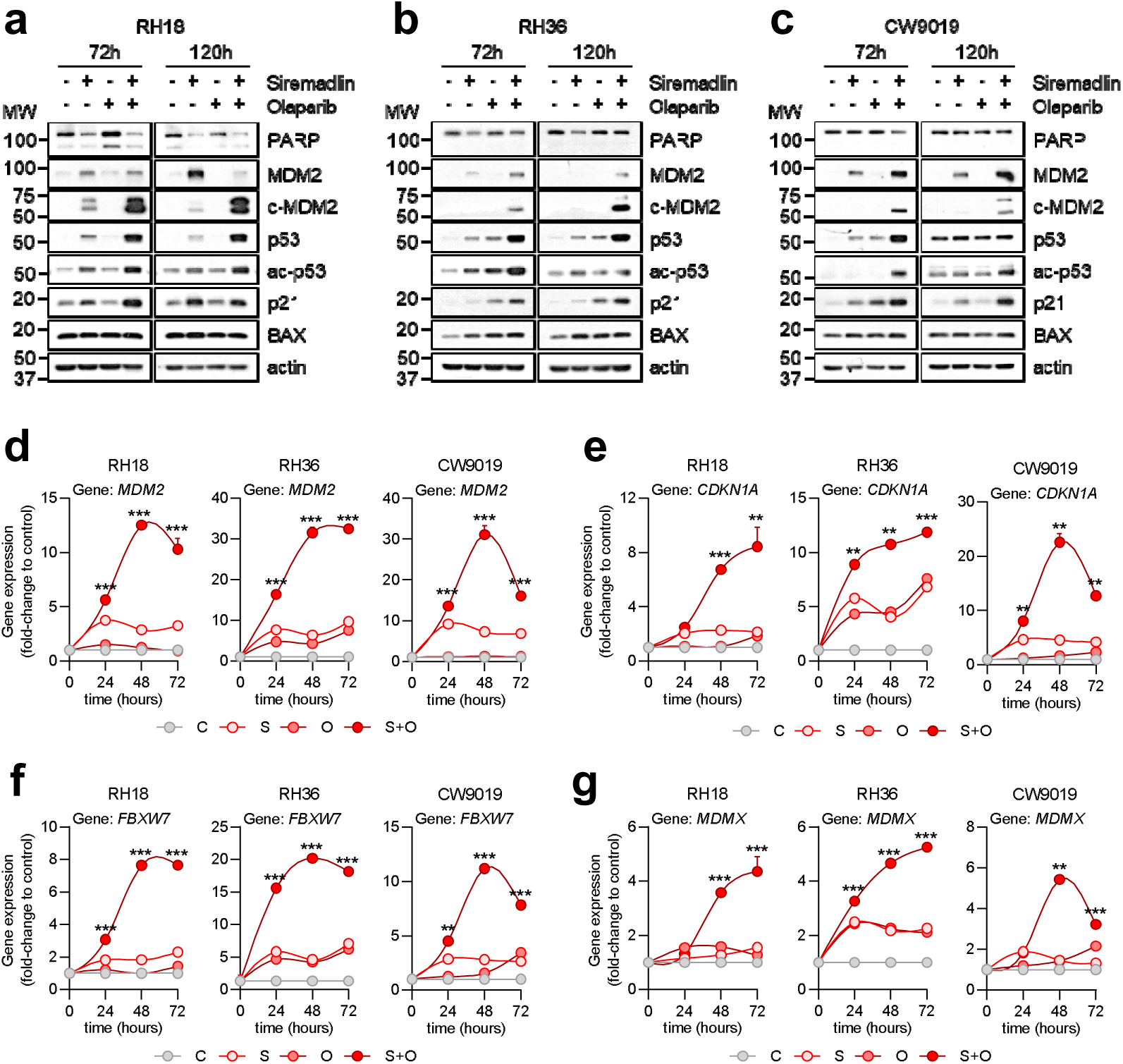
Combination of siremadlin and olaparib induces MDM2 cleavage and p53 accumulation surpassing the effects of both drugs individually. (**a** - **c**) Western blot showing the expression of PARP, full length MDM2 (MDM2), cleaved MDM2 (c-MDM2), total p53, acetylated p53 on K382 (ac-p53), p21 and BAX in p53 WT RMS cells treated with 80 nM siremadlin (S), 4 μM olaparib (O), or their combination (S+O) for 72 and 120 h. Molecular weight (MW) markers are indicated by dashes on the left side of the membrane. (**d** - **g**) Time-course plots showing the gene expression of (**d**) *MDM2*, (**e**) *CDKN1A*, (**f**) *FBXW7*, and (g) *MDMX* in individual p53 WT RMS cell lines treated with 80 nM siremadlin (S), 4 μM olaparib (O), or their combination (S+O) for 24, 48, or 72 hours, measured by qPCR. The housekeeping gene *TBP* was used as an endogenous control. Gene expression fold-change (FC) is expressed relative to the levels in vehicle-treated cells (C). Data values represent the mean ± SEM from three p53 WT RMS cell lines (*n* = 3). Statistical significance was assessed using two-way ANOVA followed by Dunnett’s post-hoc test.

To further assess p53 transcriptional activity, the expression of key p53 transcriptional targets (*MDM2*, *CDKN1A*, *FBXW7, MDMX*) was measured by qPCR at 24, 48, and 72 hours. A significant increase in *MDM2* expression was observed in p53 WT cell lines treated with siremadlin plus olaparib, starting at 24 hours and peaking at 48 hours, with a 12- to 30-fold increase compared to control (Figures 5d and S3d). Notably, the increase in *MDM2* expression was about 4- to 5-fold higher than the observed with siremadlin alone and 8- to 25-fold higher than with olaparib alone, indicating a synergistic activation of p53 activity. Consistently, similar results were obtained regarding *CDKN1A*, *FBXW7*, and *MDMX* at 48 to 72 hours post-treatment with the combination (Figures 5e-g and S3e-g). Overall, these findings demonstrate that the combination of siremadlin and olaparib enhances the individual effects of each drug by significantly boosting p53 transcriptional activity, as evidenced by the accumulation and acetylation of p53 and the increased expression of its targets, particularly MDM2 and p21.

### The combination of siremadlin and olaparib impaired tumor growth and extended mice overall survival *in vivo*

To assess the *in vivo* efficacy of the treatment with siremadlin plus olaparib, cell-derived orthotopic xenografts (CDOX) were established by injecting RH36 cells intramuscularly into the gastrocnemius of SCID mice. An initial pilot study was performed to determine an effective dose range that reduces tumor growth while minimizing toxic side effects. Hence, RH36 CDOX were treated with 50 mg/kg siremadlin, 100 mg/kg olaparib, or their combination at various concentrations (1X, 0.5X, and 0.25X) administered orally twice a week. All three combination regimens exhibited a substantial reduction in tumor growth compared to individual drug treatments as well as the control group, suggesting potential synergistic effects of the combination therapy *in vivo* (Figure 6a). However, while the combination of 50 mg/kg siremadlin + 100 mg/kg (referred to as combination 1X) yielded the most favorable outcomes, mice treated with 50 mg/kg siremadlin alone or in combination with olaparib experienced a noteworthy reduction in body weight after 30 days of administration (Figure S4a). Thus, in the subsequent *in vivo* study RH36 CDOX were treated with 25 mg/kg siremadlin and/or 50 mg/kg olaparib, with the sample size increased to six mice per group (*n* = 6). Consistently, a significant reduction in tumor growth and extended overall survival were observed in mice treated with the combination (Figure 6b-c and Figure S4b). Notably, no reduction in mice weight was observed throughout the treatments (Figure S4c). These results underscored the enhanced therapeutic efficacy of the combination regimen without causing apparent side effects at the tested doses.

**Figure 6.**
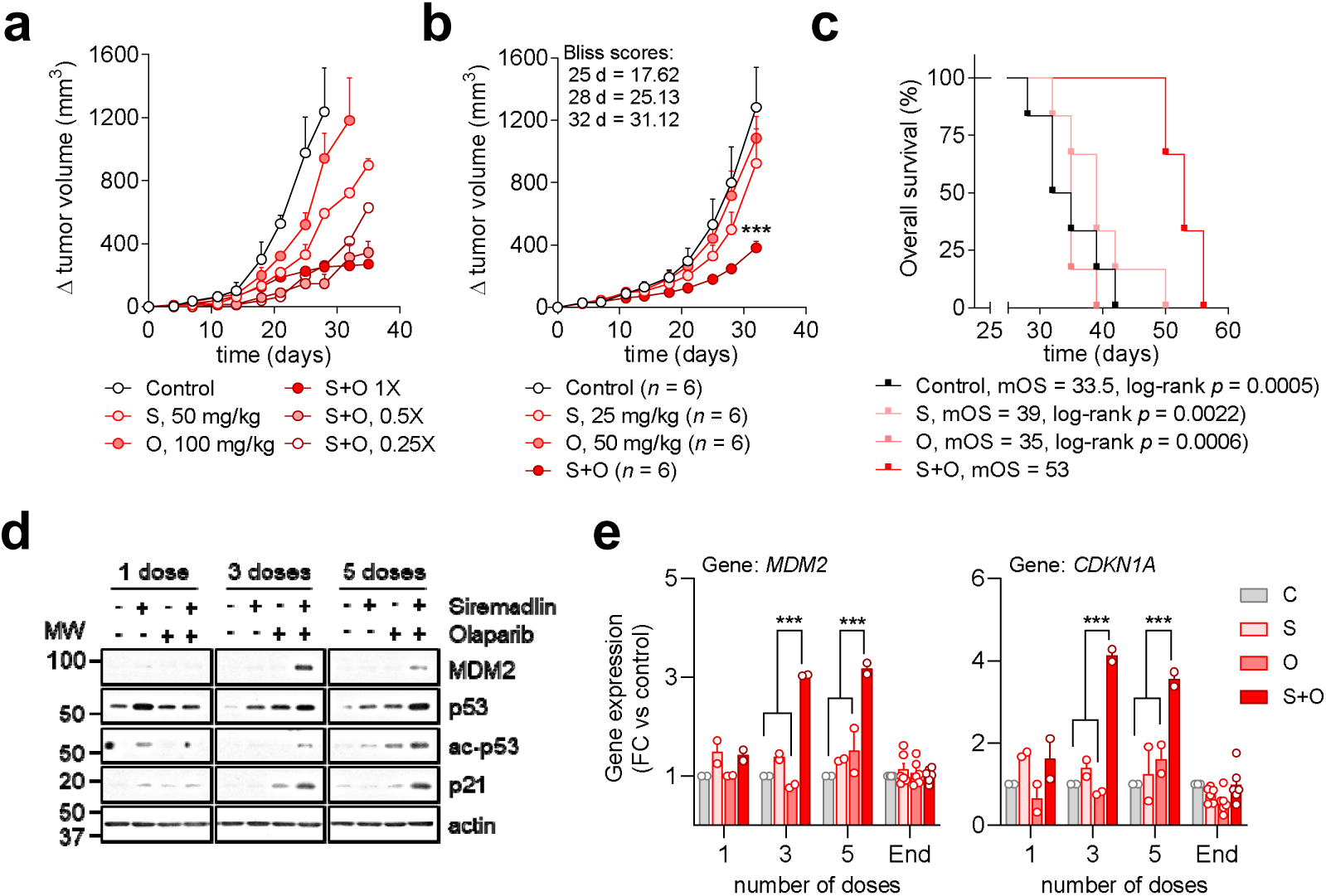
Combination of siremadlin and olaparib reduced tumor growth and extended mice overall survival. (**a**) Plot showing tumor volume increase over time for RH36-derived tumors treated with 50 mg/kg siremadlin (S), 100 mg/kg olaparib (O), or their combination at various doses: full (S+O 1X), half (S+O 0.5X), and quarter (S+O 0.25X). Each dot represents the mean ± SEM of two replicates (*n* = 2). (**b**) Plot showing the increase in tumor volume over time of RH36-derived tumors treated with 25 mg/kg siremadlin (S), 50 mg/kg olaparib (O) or their combination (S+O). Each dot represents the mean ± SEM of six tumors (*n* = 6). Two-way ANOVA with Dunnett’s post-hoc test was used to determine statistical significance. (**c**) Kaplan-Meier curves showing overall survival (in %) of mice treated with 25 mg/kg siremadlin (S), 50 mg/kg olaparib (O) or their combination (S+O). Statistical significance was assessed using log-rank test (*n* = 6). (**d**) Western blot showing the expression levels of MDM2, total p53, acetylated p53 on K382 (ac-p53), and p21 in RH36-derived tumors treated with 1, 3, or 5 doses of 25 mg/kg siremadlin (S), 50 mg/kg olaparib (O), or their combination (S+O). β-actin was used as loading control. Molecular weight (MW) markers are indicated by dashes on the left side of the membrane. (**e**) Bar plots showing the gene expression levels of *MDM2* (left) and *CDKN1A* (right) in RH36-derived tumors. Mice were treated with 25 mg/kg siremadlin (S), 50 mg/kg olaparib (O) or their combination (S+O) for 1, 3, 5 doses, or until the study endpoint (>10 doses). The housekeeping gene *TBP* was used as an endogenous control. Gene expression fold-change (mean FC ± SEM) is relative to the basal expression of each gene in vehicle-treated tumors. Statistical significance was determined using restricted maximum likelihood followed by Dunnett’s post-hoc test, accounting for the differing number of replicates across groups (*n* = 2 for tumors treated with 1, 3, or 5 doses, *n* = 6 for tumors treated until the endpoint).

Simultaneously, to assess the molecular effects of various cumulative doses, a separate cohort of RH36 CDOX received 1, 3, and 5 doses of 25 mg/kg siremadlin and/or 50 mg/kg olaparib. Protein expression analysis revealed increased levels of total and acetylated p53, as well as elevated expression of its downstream targets, p21 and MDM2, in RH36-derived tumors treated with 3 and 5 doses of the combination (Figure 6d). Additionally, gene expression analysis of some p53 targets by qPCR revealed an increased expression of *MDM2* and *CDKN1A* after 3 and 5 doses that disappeared at the endpoint (Figure 6e). Altogether, these results demonstrate the *in vivo* efficacy of combining siremadlin and olaparib, highlighting its effectiveness in reducing tumor growth and extending overall survival while enhancing p53 transcriptional activity. However, the return of *MDM2* and *CDKN1A* mRNA levels to those observed in the control and individual treatments at the study endpoint suggests an adaptive response of RH36 cells to the combination therapy, likely occurring between the fifth dose and the endpoint of the study.

### The combination of siremadlin and olaparib enhanced the therapeutic effects of selected chemotherapies

While the combination of siremadlin and olaparib showed potential in reducing tumor growth, it did not achieve complete tumor eradication as a standalone treatment. Thus, integrating siremadlin and olaparib with currently approved chemotherapy regimens for RMS was explored as a strategy to optimize clinical efficacy, minimize toxicity, and overcome treatment resistance. To assess the efficacy of combining these agents with conventional chemotherapy regimens, two distinct experimental approaches were followed (Figure 7a).

**Figure 7.**
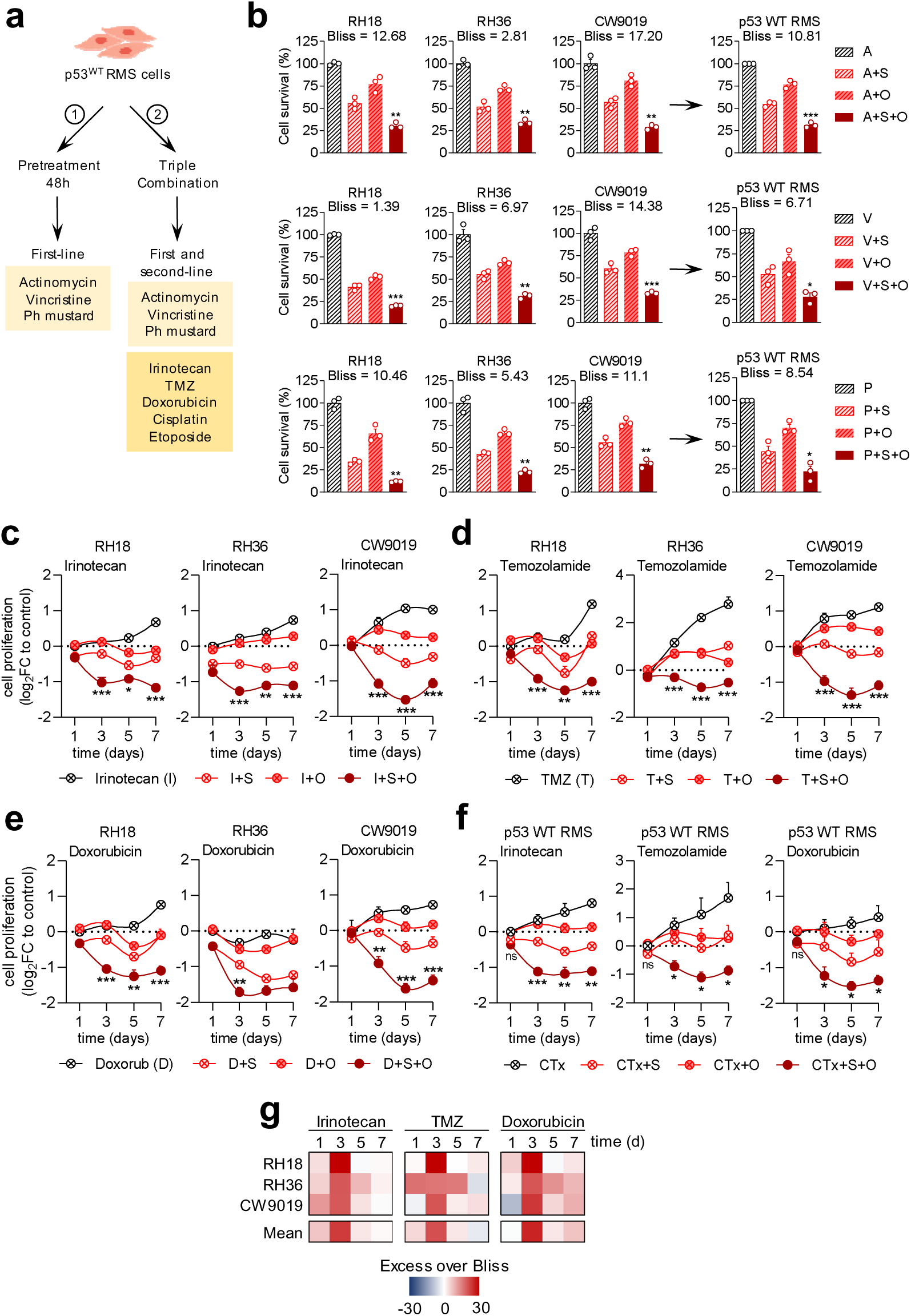
Siremadlin and olaparib retained their antitumoral efficacy following conventional chemotherapy treatments and showed synergistic effects when combined with irinotecan, temozolomide, and doxorubicin. (**a**) Schematic representation of the two distinct approaches used to evaluate the efficacy of incorporating both siremadlin and olaparib into conventional chemotherapy regimens. (**b**) Bar plots showing the effects siremadlin (S) and olaparib (O) on cell survival of p53 WT RMS cells previously treated with actinomycin-D (A), vincristine (V), or phosphoramide mustard (P) for 2 days (48 hours). Cell survival (in %) is relative to the pre-treated cells. Data represent the mean ± SEM from three replicates (*n* = 3). Statistical significance was assessed using one-way ANOVA followed by Dunnett’s post-hoc test. (**c-g**) Time-course plots showing the effects on cell proliferation in p53 WT RMS cells treated simultaneously with siremadlin (S) and olaparib (O) in combination with (**c**) irinotecan, (**d**) TMZ, or (**e**) doxorubicin. Panel (**f**) shows the mean results for these combinations across all p53 WT RMS cells. (**e**) Heatmap displaying the mean synergy Bliss scores for the combination shown in panels (c-f).

First, the efficacy of siremadlin and olaparib was evaluated in p53 WT RMS cell lines pretreated for 48 hours with vincristine, actinomycin, or phosphoramide mustard (an active cyclophosphamide metabolite). Results showed that both drugs, individually and in combination, further reduced cell survival compared to chemotherapy alone, indicating that both drugs retained their antitumoral efficacy when administered after conventional chemotherapies (Figure 7b). These findings suggest a potential additive interaction between siremadlin, olaparib, and first-line chemotherapies, leading to reduced cell survival *in vitro*.

In the second approach, p53 WT RMS cells were treated simultaneously with siremadlin and/or olaparib along with first- and second-line chemotherapeutics over a period of 7 days. Similar to the first approach, the combination of siremadlin and olaparib with first-line chemotherapeutics significantly reduced cell survival at 3 days. However, the synergistic effect diminished over time, with no statistically significant differences observed at 5 and 7 days (Figure S5a). Similar to the results with actinomycin, vincristine, and phosphoramide mustard, the combination of siremadlin and olaparib with cisplatin or etoposide produced inconsistent results (Figure S5b). Nevertheless, the combination of siremadlin and olaparib consistently demonstrated strong synergistic effects with irinotecan, temozolomide, and doxorubicin across all p53 WT RMS cell lines (Figure 7c-f). Although the synergistic effects observed at 3 days tended to become additive at 5 and 7 days (Figure 7g), these combinations continued to exert potent cytotoxic effects, significantly reducing cell proliferation compared to chemotherapy alone or dual combinations. Future research should focus on optimizing these combination strategies and elucidating the underlying mechanisms that sustain their synergistic effects. The encouraging results from combining siremadlin and olaparib with irinotecan, temozolomide, and doxorubicin present an exciting opportunity for advancing treatment protocols and improving patient care in RMS.

## Discussion

The p53-MDM2 pathway has become a prominent target for therapeutic intervention, particularly in p53 WT cancers. However, while MDM2 inhibitors were initially expected to induce strong apoptotic responses in p53 WT cells, accumulating evidence suggests that apoptosis may be constrained, likely due to specific regulatory patterns of p53 activity^33,35,36^. Furthermore, early clinical trials in advanced adult solid tumors and leukemia have shown that MDM2 inhibitors alone offer only modest therapeutic benefits, highlighting the necessity of exploring synergistic drug combinations to enhance treatment efficacy^37–39^. In RMS, preclinical studies have demonstrated encouraging results with combinations like nutlin-3 with vincristine or actinomycin D, as well as idasanutlin with radiotherapy^51,52^. However, a narrow focus on specific drug pairings may have restricted the exploration of other promising combinations involving newly developed or current therapeutic agents.

In the study presented herein, the efficacy of four promising MDM2 inhibitors (siremadlin, idasanutlin, navtemadlin, and MI-773) was evaluated as single agents in RMS. Additionally, a drug combination screening was conducted, pairing siremadlin with selected targeted therapies and standard chemotherapies commonly used in paediatric oncology. Our results demonstrated an increased vulnerability of p53 WT RMS to MDM2 inhibitors, particularly siremadlin and idasanutlin, after comparing drug sensitivities of 26 compounds between p53 WT and p53 MUT RMS cells. These findings align with previous research on first- and second-generation MDM2 inhibitors, which similarly reported differential responses to these agents in p53 WT versus p53 MUT RMS cells^51,63,64^. Notably, our study revealed a greater difference in logIC_50_ values for siremadlin compared to other inhibitors, underscoring its potential to selectively target p53 WT cells. At the molecular level, treatment with siremadlin resulted in a time-dependent accumulation of p53 and increased expression of its downstream targets, MDM2 and p21.These results are consistent with prior studies demonstrating an accumulation of p53 and subsequent upregulation of p21 in p53 WT RMS cells treated with other MDM2 inhibitors^51,63^. In addition, the induction of p21 has also been previously observed in MDM2-amplified osteosarcoma cells treated with siremadlin at various time points^65^. Collectively, our findings demonstrate an effective modulation of p53 activity through the inhibition of MDM2-p53 interaction by siremadlin in p53 WT RMS cells *in vitro*.

Given the limited clinical efficacy of siremadlin as monotherapy^37–39^, we performed a comprehensive combination drug screening involving siremadlin and 22 additional drugs in p53 WT RMS. Among the evaluated combinations, siremadlin plus olaparib exhibited the most favourable synergistic score and a substantial reduction in cellular viability, thereby emerging as the most promising treatment strategy tested. The combination of these compounds also resulted in sustained inhibition of cell proliferation over a 7-day period, exceeding the effects observed with either drug administered individually. Moreover, cell painting analysis revealed a distinct clustering of cells treated with both drugs, indicating unique phenotypic effects compared to individual treatments. This distinct clustering strongly supports the hypothesis of a potential synergistic interaction between the compounds, significantly influencing multiple cellular processes and resulting in a novel cellular phenotype. These findings are consistent with a previous study that demonstrated synergistic effects when combining idasanutlin with rucaparib (a PARP inhibitor) in both PARP-sensitive and PARP-resistant ovarian cancer cell lines, suggesting that MDM2-p53 inhibition may resensitize cells to PARP inhibition^66^.

Both MDM2 and PARP inhibitors are known to induce p53 activation and p21-mediated cell cycle arrest^67–69^. However, the individual efficacy of these compounds is partly constrained by the challenge of transitioning from pulsatile to sustained p53 activation^36,65,70^. Pulsatile p53 activation involves the induction of proteins that mediate cell cycle arrest, such as p21, and proteins that maintain negative feedback loops crucial for generating p53 activation pulses, like MDM2^70–73^. In contrast, sustained p53 activity can drive intrinsic apoptosis through various mechanisms, particularly in cells that are unable to repair DNA damage or are overexposed to excessive stress signals^36,71,73,74^. In our study, the combination of siremadlin and olaparib resulted in a significant increase in the number of apoptotic and dead p53 WT RMS cells, surpassing the effects observed with the single treatments. At the same time, we also observed a decreased expression of p-Rb and cyclin B1 that supported the impaired cell cycle progression observed in p53 WT cells treated with both drugs. At the molecular level, the combination of siremadlin and olaparib significantly increased MDM2 cleavage in all p53 WT RMS cells. This finding is particularly relevant because the ∼60 kDa MDM2 species (MDM2-p60), produced by caspase-2-PIDDosome and/or caspase-3 cleavage, stabilize p53 and shift its dynamics from pulsatile to sustained mode, thereby promoting apoptosis in extensively damaged cells^36,75,76^. Indeed, concomitant with the observed MDM2 cleavage, our results revealed a strong increase in both total and acetylated p53 (ac-p53) levels. The acetylation of p53 at lysine 382 is also significant, as this modification enhances the transcriptional activity of p53^77,78^. Consistent with the elevated levels of p53 expression and acetylation, a strong transcriptional upregulation of p53 target genes is observed in response to the combination therapy, significantly exceeding the induction levels seen with the single treatments. The increased induction of p21 expression compared to single treatments is also relevant, as p21 is a pivotal mediator of p53-induced cell cycle arrest. This result substantiates the previously observed alterations in cell cycle progression, shedding light on the molecular mechanisms by which the combination therapy exerts its cytotoxic effects. Overall, these findings suggest that combining siremadlin and olaparib significantly enhances the efficacy of PARP inhibitors by disrupting the MDM2-p53 negative feedback loop following initial p53 activation, leading to sustained p53 activation. Additionally, the transition from pulsatile to sustained p53 activation appears to be further supported by MDM2 cleavage induced by the combination treatment. The resulting increase in p53 stabilization and sustained activation likely contributes to the pronounced apoptotic response observed. These findings highlight the potential of combining siremadlin and olaparib as a potent therapeutic strategy, overcoming the limitations of each drug when used individually and offering a more effective approach for cancer treatment.

The *in vivo* evaluation of the combination of siremadlin and olaparib in RH36 xenografts provided valuable insights into both the therapeutic potential and the underlying molecular mechanisms of this treatment. The dosing regimen for siremadlin was informed by prior research, which demonstrated that pulsed high-doses of siremadlin induce a stronger proapoptotic response in p53 WT cancer cells compared to sustained low-doses^65^. For olaparib, the dosing regimen was based on established efficacy from previous studies, which identified effective oral doses of 50 or 100 mg/kg daily in xenograft models^79–81^. In this study, the administration intervals for olaparib were adjusted to a biweekly schedule to align with the dosing regimen of siremadlin. Remarkably, the combination 25 mg/kg siremadlin and/or 50 mg/kg olaparib regimen significantly reduced tumor growth and extended overall survival in mice compared to individual treatments and control groups, demonstrating synergistic therapeutic efficacy with no apparent side effects. At the molecular level, the combination treatment led to an increase in both total and acetylated p53, as well as elevated protein and mRNA levels of its downstream targets, p21 and MDM2. These findings provide evidence that the combination of siremadlin and olaparib significantly enhanced p53 transcriptional activity *in vivo*, compared to individual treatments and control.

Finally, the integration of siremadlin plus olaparib after the treatment with first-line chemotherapies or in combination with particular chemotherapies such as irinotecan, doxorubicin or temozolomide showed additive/synergistic effects, suggesting a potential strategy for enhancing the therapeutic outcomes of second-line treatments for RMS while reducing the dose-limiting toxicities and adverse effects associated with these therapies.

Overall, this study presents the first compelling evidence demonstrating the enhanced therapeutic efficacy of combining siremadlin and olaparib in the treatment of p53 WT RMS *in vivo*. The substantial reduction in tumor growth and extended survival, along with molecular evidence of p53 pathway activation, underscores the strong potential of this combination therapy *in vivo*. The strong additive and/or synergistic effects observed with irinotecan, temozolomide, and doxorubicin provide a compelling rationale for further preclinical optimization of dose regimens and the exploration of these combinations in both preclinical and clinical settings.

## Conclusion

Cancer cells often exhibit elevated DNA damage and replication stress, making them more dependent on PARP. While PARP inhibitors have demonstrated efficacy in homologous recombination-deficient tumors, their success in other cancer types has been limited. Our study demonstrates that combining siremadlin with olaparib effectively exploits cancer cells’ vulnerability to PARP inhibition while disrupting MDM2-mediated negative feedback. This combined approach not only prolongs and amplifies the p53 response, activating cell death pathways in p53 WT RMS cells both *in vitro* and *in vivo*, but it also allows for dose reduction of each drug, which may lead to decreased treatment-related side effects. These findings provide a compelling rationale for further clinical exploration of siremadlin and olaparib in treating p53 WT RMS and offer valuable insights into novel multidrug therapeutic strategies that may also be applicable to other solid tumors with similar molecular features, potentially improving patient outcomes with fewer adverse effects.

## Supporting information

Supplementary Table S1. List of cell lines

Supplementary Table S2. Number of cells seeded

Supplementary Table S3. Drugs and doses used

Supplementary Table S4. List of antibodies used for WB

## Acknowledgments

The authors wish to thank the members of the High Technology Unit (UAT) and the Laboratory Animal Service (LAS) at the Vall d’Hebron Research Institute for their scientific and technical support. This work was funded by the *Instituto de Salud Carlos III* through the projects [PI21/00640] and [FORT23/00034] (co-financed by the European Regional Development Fund (ERDF) under the program “A way to make Europe”). Additional financial support was provided by the Catalan Institute of Oncology (ICO), FEDER Funds, AGAUR (grant 2021 FI_B 00088), *Fundació A. BOSCH*, *Amics Joan Petit*, *Fundación Inocente*, *Iniciativa Tot per Tu*, and *Mi Compañero de Viaje*.

## Author’s contributions

Conceptualization: G.P., L.M. and J.R.; methodology: G.P., P.Z., G.G.-O., A.W., and M.L; formal analysis: G.P. and J.R.; data curation: G.P., A.W., M.L., and J.C.-P.; writing—original draft preparation: G.P., J.C.-P., L.M. and J.R.; writing—review and editing: G.P., P.Z., G.G.-O, L.G.-G., J.S.-G., N.N., M.F.S., A.S., G.G., R.H., J.C.-P., L.M. and J.R; supervision: L.M. and J.R.; project administration: J.R.; funding acquisition: J.S.d.T., S.G., L.M. and J.R. All authors have read and agreed to the published version of the manuscript.

