## Supplementary Figures S1-S5 for "Combined inhibition of MDM2 and PARP lead to a synergistic anti-tumoral response in p53 wild-type rhabdomyosarcoma models"

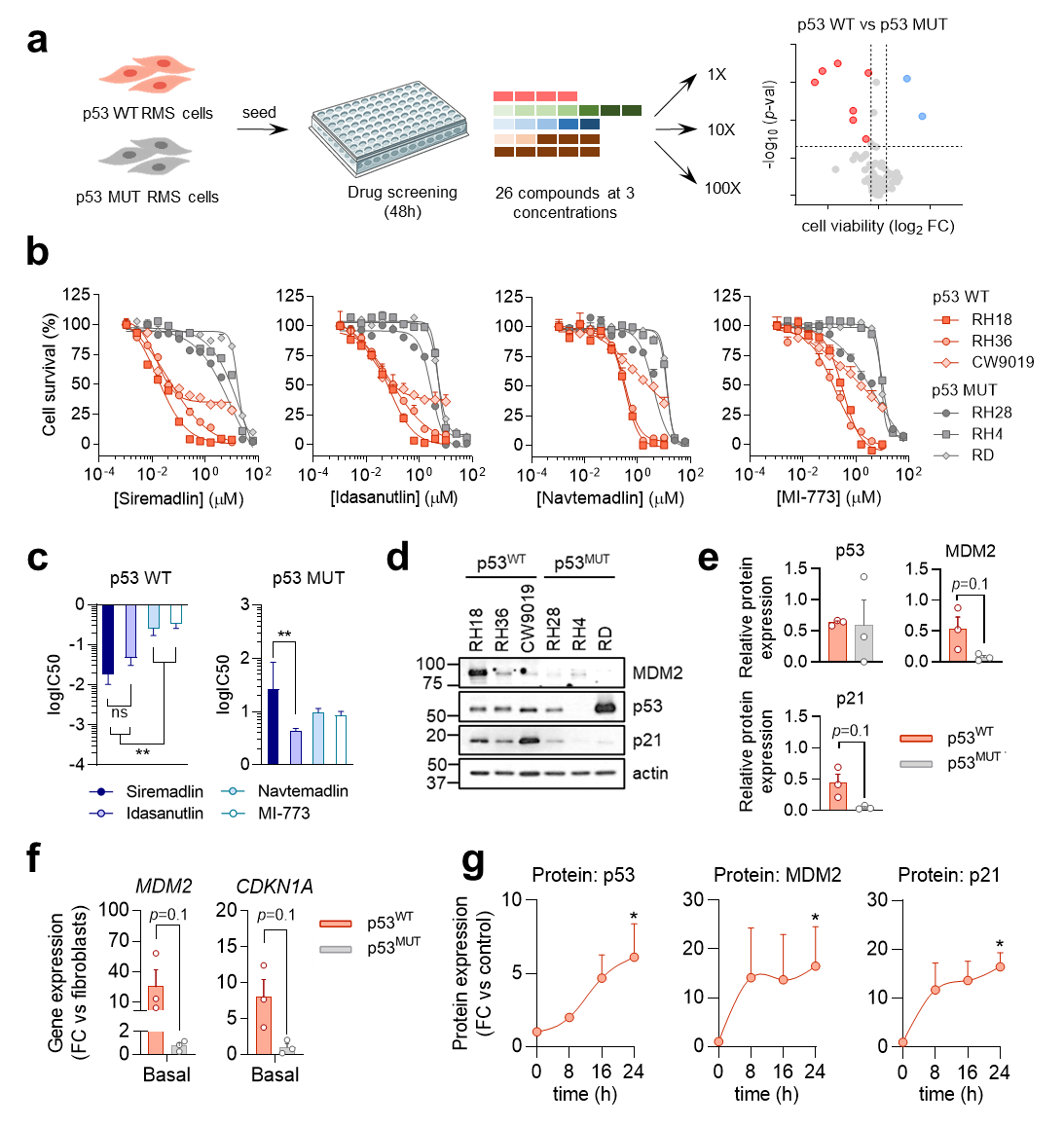
**Supplementary Figures**

**Supplementary Figure S1. Comprehensive assessment of MDM2 inhibitors in p53 WT and p53 MUT RMS cells.** (**a**) Scheme of the initial drug screening workflow conducted in both p53 WT and p53 MUT RMS cells. (**b**) Dose-response curves for siremadlin, idasanutlin, navtemadlin, and MI-733 (from left to right) across the six cell lines used. IC_50_ values were determined using a non-linear regression approach (least squares regression without weighting). Dots represent the mean ± SEM of three technical replicates (*n* = 3). (**c**) Comparative analysis of logIC_50_ values in p53 WT (left) and p53 MUT (right) highlighting the relative potency of MDM2 inhibitors in each cellular context. Bars represent the mean ± SEM of three p53 WT and three p53 MUT RMS cells (*n* = 3). Statistical significance was assessed using one-way ANOVA followed by Tukey’s post-*hoc* test. (**d**) Western blot showing the basal expression of MDM2, p53 and p21 across six RMS cells. β-actin was used as the loading control. (**e**) Bar plots displaying the expression levels of MDM2, p53, and p21 relative to β-actin. Data are presented as the mean ± SEM of three p53 WT and three p53 MUT RMS cells (*n* = 3). Statistical significance between both groups was assessed using the non-parametric Mann Whitney’s test. (**f**) Bar plots showing the basal gene expression of *MDM2* and *CDKN1A* in three p53 WT RMS cells and three p53 MUT RMS cells (*n* = 3). The housekeeping gene *TBP* was used as endogenous control. Gene expression fold-change (mean ± SEM) is relative to the basal expression of fibroblasts. Statistical significance between both groups was assessed using the non-parametric Mann Whitney’s test. (**g**) Bar plots showing the protein quantification normalized to β-actin of results presented in Figure 1d. Data represent the mean ± SEM from three p53 WT RMS cells (*n* = 3). Statistical significance was assessed using one-way ANOVA followed by Kruskal-Wallis post-hoc test.

**Supplementary Figure S2. Phenotype profiling revealed dose-dependent morphologic changes induced by siremadlin and olaparib in RH36 cells.** (**a**) High-content images from cell painting analysis of RH36 cells treated with vehicle, 80 nM siremadlin, 3.2 μM olaparib, or their combination. The merged images display Hoechst-stained nuclei in blue, mitochondria in red, and the cytoskeleton and Golgi apparatus in green. (**b**) Two-dimensional plot, derived from *t*-SNE analysis, representing the morphological changes triggered by siremadlin and olaparib in comparison to other compounds in RH36 cells. Dot colors represent different treatments, and dot sizes indicate the doses of the compound tested, with larger dots corresponding to higher doses. (**c**) Heatmap showing the hierarchical clustering of >2000 morphologic features analyzed by cell painting after treating RH36 cells with increasing doses of siremadlin and olaparib. The color in the heatmap (Z-score) represents the deviation of each morphological feature from the mean value of the DMSO-treated control cells. (**d**) On the left, radial plots show the sum of both positive and negative differences, while on the right results are categorized in positive and negative differences. Concentric marks indicate Z-scores. All morphological features analyzed are grouped into eight main groups: endoplasmic reticulum (ER), nucleus, nucleoli, Golgi + cytoskeleton, mitochondria, cellular area and shape, characteristics of the neighbor cells, and correlates between nucleus and cytoplasm regions. At the same time, these features are further categorized into the following subgroups: RD (radial distribution), L (location and spatial distribution), G (granularity), I (integrated intensity), C (cell measurements), N (nuclear measurements), and Cy (cytoplasm measurements).

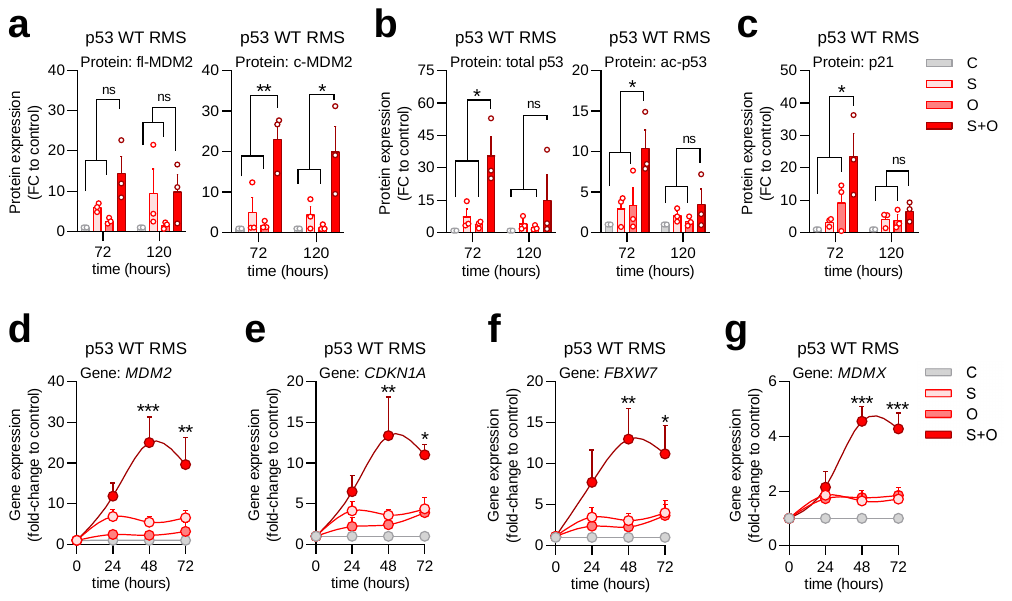

**Supplementary Figure S3. Quantification of protein and gene expression in p53 WT RMS cells treated with siremadlin and olaparib, alone or in combination.** (**a-c**) Bar plot showing the quantification of blots shown in Fig. 5a-c for (**a**) full length and cleaved MDM2 (c-MDM2), (**b**) total and acetylated p53, and (**c**) p21 in p53 WT RMS cells treated with 80 nM siremadlin (S), 4 μM olaparib (O), or their combination (S+O) for 72 and 120 hours. Protein expression was normalized to β-actin and is expressed in fold-change (FC) relative to vehicle-treated cells (C). Data values are expressed as the mean ± SEM of three p53 WT RMS cells (*n* = 3). Statistical significance was assessed using two-way ANOVA with Dunnett’s post-hoc test. (**d-g**) Dot plots showing the mean gene expression of (**d**) *MDM2*, (**e**) *CDKN1A*, (**f**) *FBXW7*, and (**g**) *MDMX* calculated from the three p53 WT RMS cells shown in Figure 4d-g. The housekeeping gene *TBP* was used as an endogenous control. Gene expression fold-change (FC) is expressed relative to the levels in vehicle-treated cells (C). Data values represent the mean ± SEM from three p53 WT RMS cells (*n* = 3). Statistical significance was assessed using two-way ANOVA followed by Dunnett’s post-hoc test.

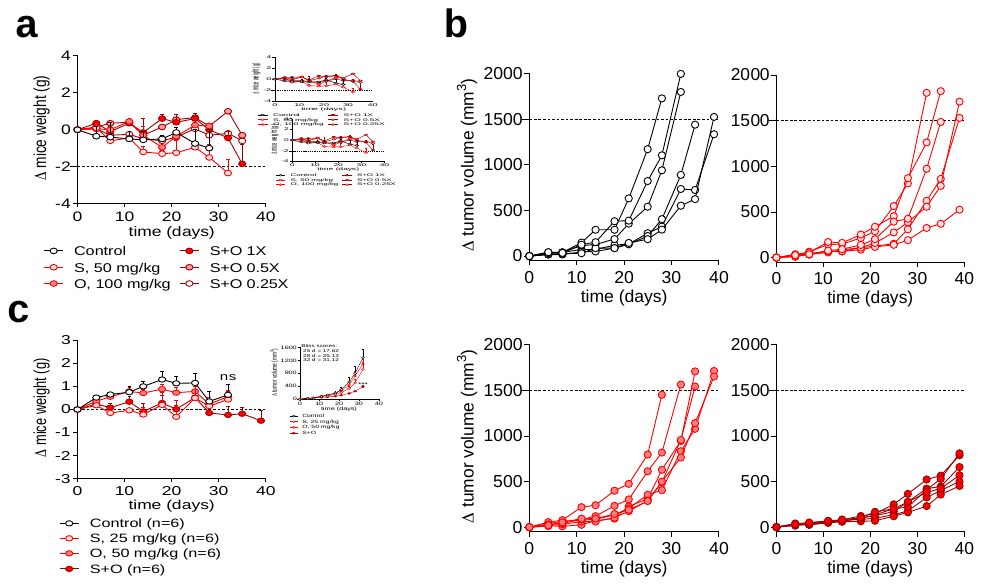

**Supplementary Figure S4. *In vivo* efficacy and dose accumulation effects of the siremadlin and olaparib combination in RH36-derived tumors.** (**a**) Dot plot showing the change in mouse weight over time for mice treated with 50 mg/kg siremadlin (S), 100 mg/kg olaparib (O), or their combination at full doses (S+O 1X), half doses (S+O 0.5X), and quarter doses (S+O 0.25X). Each dot represents the mean ± SEM from two replicates (*n* = 2). (**b**) Dot plot showing the increase in tumor volume over time of RH36-derived tumors treated with 25 mg/kg siremadlin (S), 50 mg/kg olaparib (O), or their combination (S+O). Each dot represents an individual tumor. (**c**) Dot plot representing the change in weight over time of mice treated with 25 mg/kg siremadlin (S), 50 mg/kg olaparib (O), or their combination. Each dot represents the mean ± SEM of six RH36-derived tumors (*n* = 6). Two-way ANOVA with Dunnet’s post-*hoc* test was used to determine statistical significance.

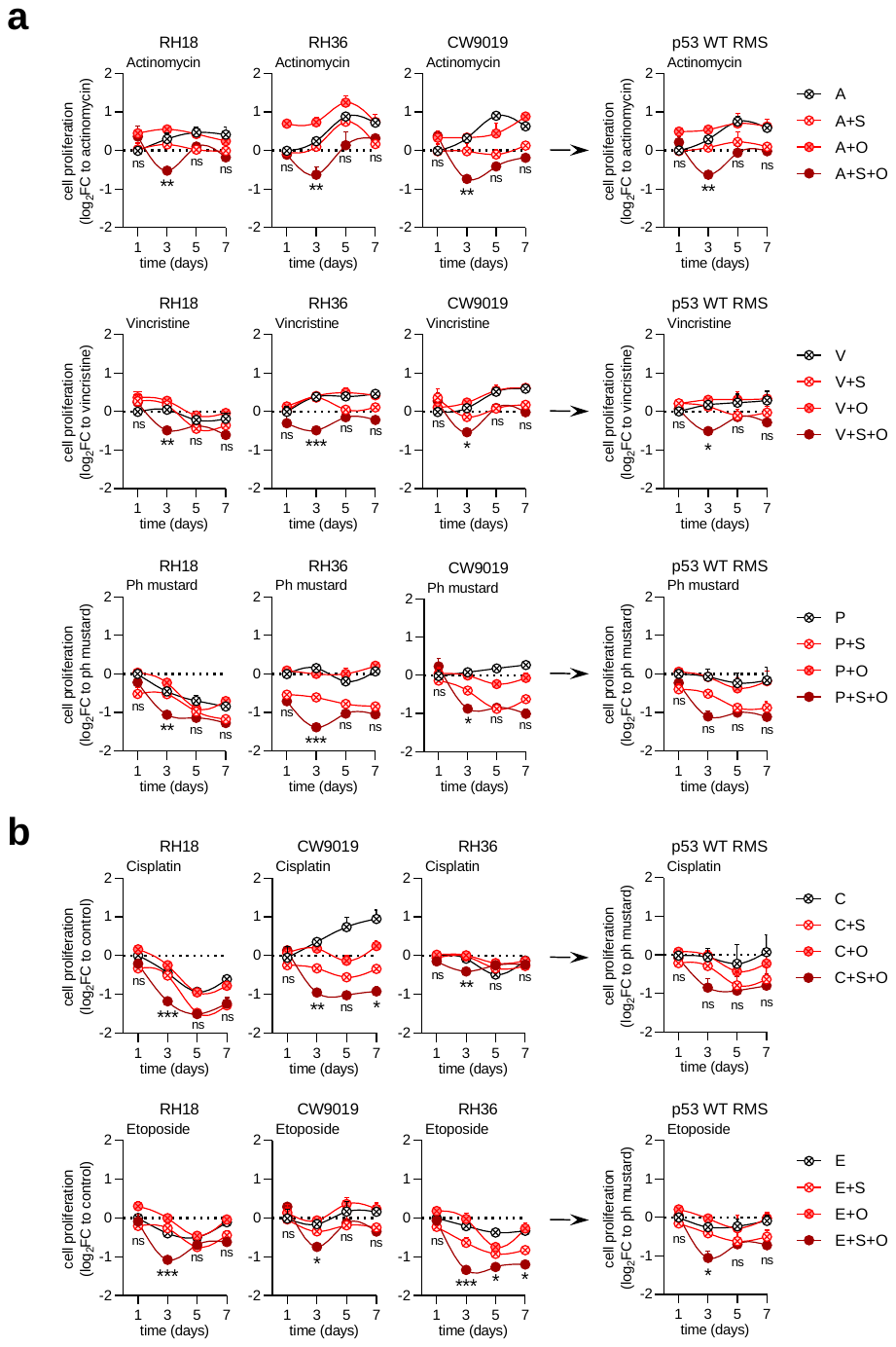

**Supplementary Figure S5. Effect of combining siremadlin and olaparib with chemotherapeutics on p53 WT RMS cell proliferation.** (**a-b**) Dot plots showing the effect on cell proliferation of combining siremadlin (S) and olaparib (O) with (**a**) first-line chemotherapeutics and (**b**) cisplatin or etoposide. Cell proliferation is expressed as log_2_ fold-change (log_2_FC) relative to cells treated with chemotherapy alone. In all cases, cell proliferation is expressed as log_2_ fold-change (log_2_FC) relative to cells treated with chemotherapy alone. Data represent the mean ± SEM from three replicates (*n* = 3). Statistical significance was assessed using two-way ANOVA followed by Dunnett’s post-hoc test.
